# EGFR confers exquisite specificity of Wnt9a-Fzd9b signaling in hematopoietic stem cell development

**DOI:** 10.1101/387043

**Authors:** Stephanie Grainger, Nicole Nguyen, Jenna Richter, Jordan Setayesh, Brianna Lonquich, Chet Huan Oon, Jacob M. Wozniak, Rocio Barahona, Caramai N. Kamei, Jack Houston, Marvic Carrillo-Terrazas, Iain A. Drummond, David Gonzalez, Karl Willert, David Traver

## Abstract

The mechanisms of Wnt-Frizzled (Fzd) signaling selectivity and their biological implications remain unclear. We demonstrate for the first time that the epidermal growth factor receptor (EGFR) is required as a co-factor for Wnt signaling. Using genetic studies in zebrafish, paired with *in vitro* cell biology and biochemistry, we have determined that Fzd9b signals specifically with Wnt9a *in vivo* and *in vitro* to elicit β-catenin dependent Wnt signals that regulate hematopoietic stem and progenitor cell (HSPC) development in the dorsal aorta. This requirement is conserved in the derivation of HSPCs from human embryonic stem cells. Wnt9a-Fzd9b specificity requires two intracellular domains in Fzd9b, which interact with EGFR as a required co-factor to promote signal transduction. EGFR phosphorylates one tyrosine residue on Fzd9b, a requirement for the Wnt signal. These findings indicate that Wnt signaling interactions can be exquisitely specific and inform protocols for derivation of HSPCs *in vitro*.

**Highlights:**

- An *in vitro* signaling screen identifies Fzd9b as a Wnt9a-specific receptor.
- Fzd9b and Wnt9a regulate hematopoietic stem cell development as a cognate pair.
- WNT9A and FZD9 are required for HSPC derivation from human pluripotent cells *in vitro*.
- EGFR confers specificity to Wnt9a-Fzd9b signaling in zebrafish and human cells.

## Introduction

Members of the *Wnt* gene family encode highly conserved, lipid-modified glycoproteins that are involved in the regulation of a plethora of developmental processes including stem cell maintenance, proliferation and differentiation, as well as embryonic patterning and morphogenesis (reviewed in Loh et al., 2016). Although the mammalian genome encodes 19 *Wnts* and 10 *Frizzled* (*Fzd*) receptors, there is little evidence for signaling specificity through cognate Wnt-Fzd pairings (Cho et al., 2017). Previously, we established that Wnt9a is required specifically in zebrafish and human hematopoiesis, in that several other Wnt ligands could not replace the loss of Wnt9a (Grainger et al., 2016b; Richter et al., 2018). Using hematopoietic stem and progenitor cell (HSPC) development as a platform for validation, we demonstrate for the first time that the epidermal growth factor receptor (EGFR) is required as a co-factor to mediate the exquisite specificity of the WNT9A-FZD9 signaling interaction, a finding which may indicate a general paradigm for regulating Wnt-Fzd signaling specificity.

Hematopoietic stem cells (HSCs) are the tissue-specific stem cells that provide blood and immune cells for the duration of an organism’s life. During development, these cells arise directly from major arterial vessels, from a specialized population of cells termed hemogenic endothelium (HE) (Bertrand et al., 2010; Cumano et al., 2001; Jaffredo et al., 1998; Kissa and Herbomel, 2010; Medvinsky and Dzierzak, 1996; Zovein et al., 2008). Prior to their emergence from the vasculature, cells of the HE are specified from mesodermal progenitors in the lateral plate mesoderm (Brown et al., 2000; Fouquet et al., 1997; Herbert et al., 2009; Jin et al., 2005; Liao et al., 1997). As these cells ingress beneath the somites, they receive inductive cues from developmental regulators, including fibroblast growth factors (FGF), Notch, and Wnt, to establish their fate and future function as aorta, vein, or HE from nearby tissues such as the somite and neural crest cells (Bertrand et al., 2010; Burns et al., 2005; Butko et al., 2015; Clements et al., 2011; Clements and Traver, 2013; Damm and Clements, 2017; Kobayashi et al., 2014; Leung et al., 2013; Wilkinson et al., 2009; Zhen et al., 2013). After their specification, HSPCs emerge directly from ventral endothelium comprising the dorsal aorta (hereafter aorta) in a process termed the endothelial-to-hematopoietic transition (EHT) (Bertrand et al., 2010; Kissa and Herbomel, 2010). They then enter circulation and migrate to secondary hematopoietic organs such as the fetal liver in mammals, or the caudal hematopoietic tissue in teleosts, where they are thought to amplify and mature (Murayama et al., 2006; Tamplin et al., 2015) before seeding the final sites of hematopoietic residence in the bone marrow of mammals, or the kidney marrow of teleosts (Jagannathan-Bogdan and Zon, 2013).

Among the multiple inductive signals regulating HSC development and homeostasis are the growth factors encoded by the *Wnt* gene family (Baba et al., 2006; Fleming et al., 2008; Goessling et al., 2009; Kirstetter et al., 2006; Luis et al., 2012; Luis et al., 2011; Luis et al., 2009; Malhotra et al., 2008; Reya et al., 2003; Scheller et al., 2006; Willert et al., 2003; Zhao et al., 2007). We previously determined that an early Wnt9a cue drives a proliferative event in the aorta, after HSCs have emerged, but before they have seeded the secondary hematopoietic organs. Interestingly, this process cannot be mediated by other Wnt ligands, including Wnt3a or Wnt9b, suggesting exquisite specificity of Wnt ligand function, which may be mediated through specific interaction with one of the 14 zebrafish Fzd receptors (Grainger et al., 2016b).

Here, we identify Fzd9b as the cognate signaling partner for Wnt9a in the process of hematopoietic stem and progenitor cell (HSPC) development. Wnt9a and Fzd9b loss of function phenotypes in hematopoietic development are indistinguishable, and genetic complementation indicates that each operates in the same genetic pathway, upstream of β-catenin. This ligand-receptor pairing is conserved in human hematopoiesis *in vitro*, as determined using a previously established protocol to generate HSPCs (Ng et al., 2008). A series of chimeric receptors between Fzd9b and a Fzd that does not promote Wnt9a signaling reveals that intracellular Fzd9b domains mediate the specificity of this Wnt-Fzd pairing, both *in vitro* and *in vivo*, indicating that a transmembrane spanning co-factor is involved in establishing specificity. Using a proximity ligation method followed by mass spectrometry (Hung et al., 2016; Lam et al., 2015), we identified the receptor tyrosine kinase EGFR as required for this specific signaling interaction for both zebrafish and human proteins. Altogether, these results demonstrate a conserved Wnt-Fzd pairing that mediates a precise Wnt cue required for hematopoiesis and opens the field for discovery of other Wnt-Fzd specificity co-regulators.

## Results

### Fzd9b interacts with Wnt9a *in vitro*

We had previously established a specific requirement for Wnt9a in directing an early amplification of HSPCs, a surprising finding because Wnts have been thought to be functionally promiscuous (Grainger et al., 2016b; Ring et al., 2014; Voloshanenko et al., 2017). Vertebrate genomes encode multiple Fzd receptors, with 10 *Fzd* genes in mammals and 14 *fzd* genes in zebrafish, suggesting that one mechanism for specific Wnt function involves specific Wnt-Fzd pairing. Wnt9a signals from the somite to ingressing HE prior to 20 hours post fertilization (hpf) (Grainger et al., 2016b). A screen for *fzd* expression in 16.5 hpf *fli1a*-positive cells indicated that a majority of Fzds were expressed in the ingressing endothelial cell population (Fig. S1A-B). To narrow down the number of potential Wnt9a receptors, we employed an established β-catenin dependent Wnt reporter assay (Veeman et al., 2003), called Super-TOP-Flash (STF). Wnt9a alone induced low levels of STF reporter activity; however, upon co-transfection with Fzd5 and Fzd9b, but no other Fzd, Wnt9a significantly increased STF reporter activity (Fig. S1C-D, Fig. 1A). Since we hypothesized that Wnt9a would signal to its cognate receptor on neighboring cells, we used a co-culture approach to assess the ability of Wnt9a to act in a paracrine manner on Fzd5 and Fzd9b. In this co-culture assay, Fzd9b, but not Fzd5, was able to transduce the Wnt9a signal and activate STF reporter activity (Fig. 1B-C), indicating that Fzd9b acts as a specific Wnt9a receptor. Taken together, these results indicate that Fzd9b is able to transduce the Wnt9a signal *in vitro,* suggesting that Fzd9b is involved in Wnt9a-mediated HSC development.

**Figure 1:**
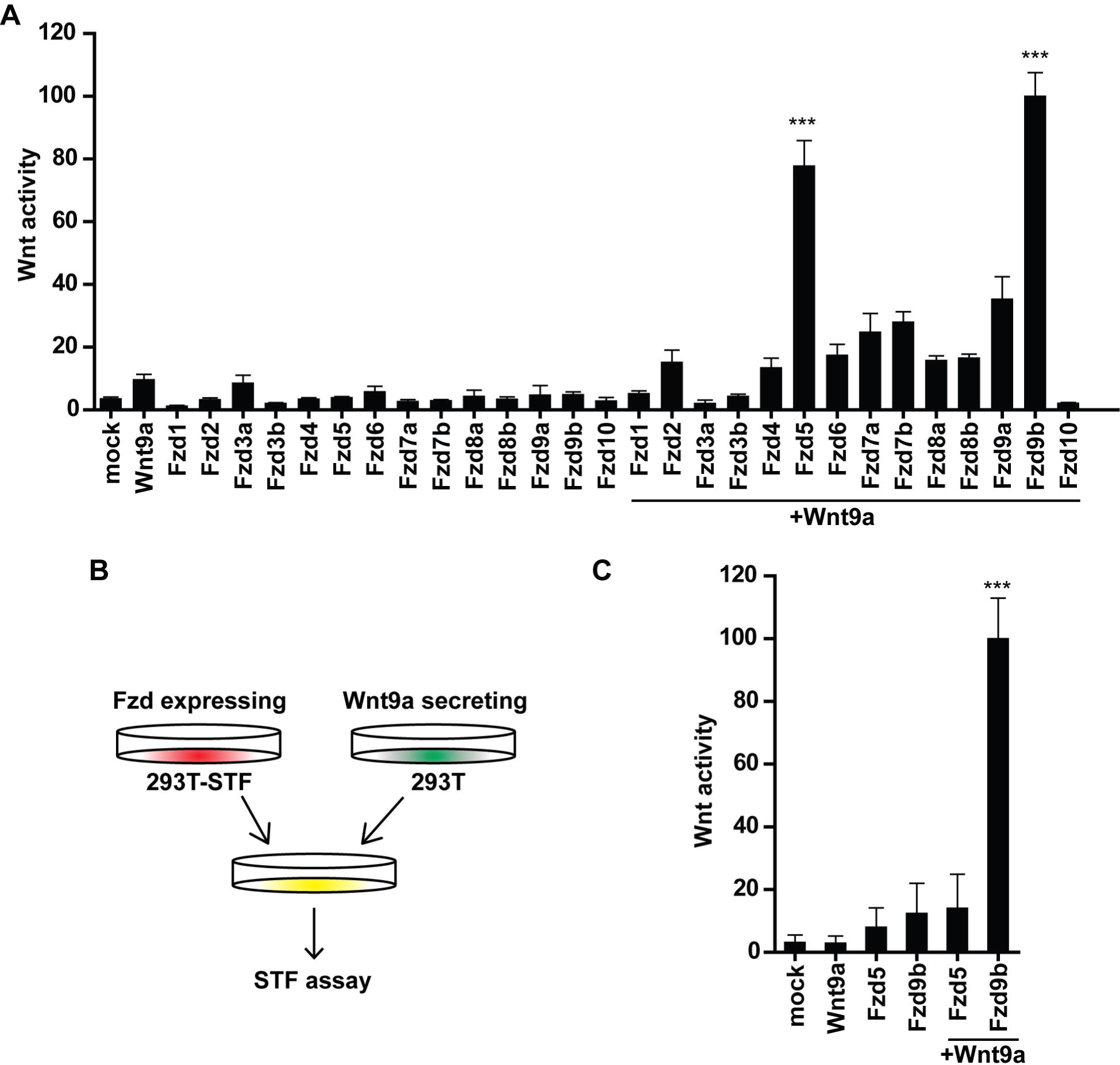
Fzd9b interacts with Wnt9a *in vitro*. **A.** Quantification of HEK293T cell STF assay screen with zebrafish Wnt9a and Fzds. Bars represent the mean of a minimum of 3 biological replicates; error bars represent standard deviation; ***P<0.001 compared to Wnt9a alone. **B.** Co-culture assay set up for **C. C.** HEK293T cells expressing zebrafish Wnt9a were mixed with HEK293T STF cells expressing zebrafish Fzd9b or Fzd5 and assayed for Wnt activity. Bars represent the mean of a minimum of 3 biological replicates; error bars represent standard deviation; ***P<0.001 compared to Wnt9a cells with mock transfected HEK293T STF cells. (See also Figure S1).

### Fzd9b is required for zebrafish hematopoiesis

Since somite-derived Wnt9a signals to ingressing HE prior to 20 hpf, we hypothesized that *fzd9b* would be expressed in cells of the HE prior to this. By whole-mount *in situ* hybridization (WISH) or double-fluorescent whole-mount *in situ* hybridization (FISH) at 15 hpf, we detected *fzd9b* in neural tissues, in the anterior lateral plate mesoderm, which contains endothelial cells contributing mostly to primitive hematopoiesis (Detrich et al., 1995; Holz et al., 2003; Zhang and Rodaway, 2007), and in the medial and posterior lateral plate mesoderm, the tissues from which HE is derived (Maeno et al., 1985; Pardanaud et al., 1996; Turpen et al., 1981) (Fig. 2A). FISH for *fzd9b* and the endothelial marker *fli1a* confirmed that *fzd9b* is expressed in endothelial precursor cells (Fig. 2B). To test if hematopoietic cells were derived from cells expressing *fzd9b*, we performed a lineage tracing experiment using *fzd9b* promoter sequences driving expression of Gal4 in two ways. First, Gal4 can activate an *upstream activating sequence* (UAS) driven GFP (*fzd9b:Gal4;UAS:GFP);* secondly *UAS:Cre* can be similarly activated, which ultimately leads to recombination of loxP sites flanking a BFP, and expression of dsRed (*fzd9b:Gal4; UAS:Cre; loxP BFP loxP dsRed)* Using this strategy, with which we were able to observe GFP-positive or dsRed-positive (pseudo colored as green here) cells in the floor of the dorsal aorta in the characteristic cup shape seen during the EHT (at 40 hpf), indicating that nascent HSPCs had expressed *fzd9b* prior to their emergence (Fig. 2C, Fig. S2A). T-cells derived from HSCs reside in the thymus beginning around 4 days post-fertilization (dpf) and expressed GFP at 6 and 7 dpf (Fig. 2D, Fig S2B), consistent with a function for *fzd9b* in HSPC development.

**Figure 2:**
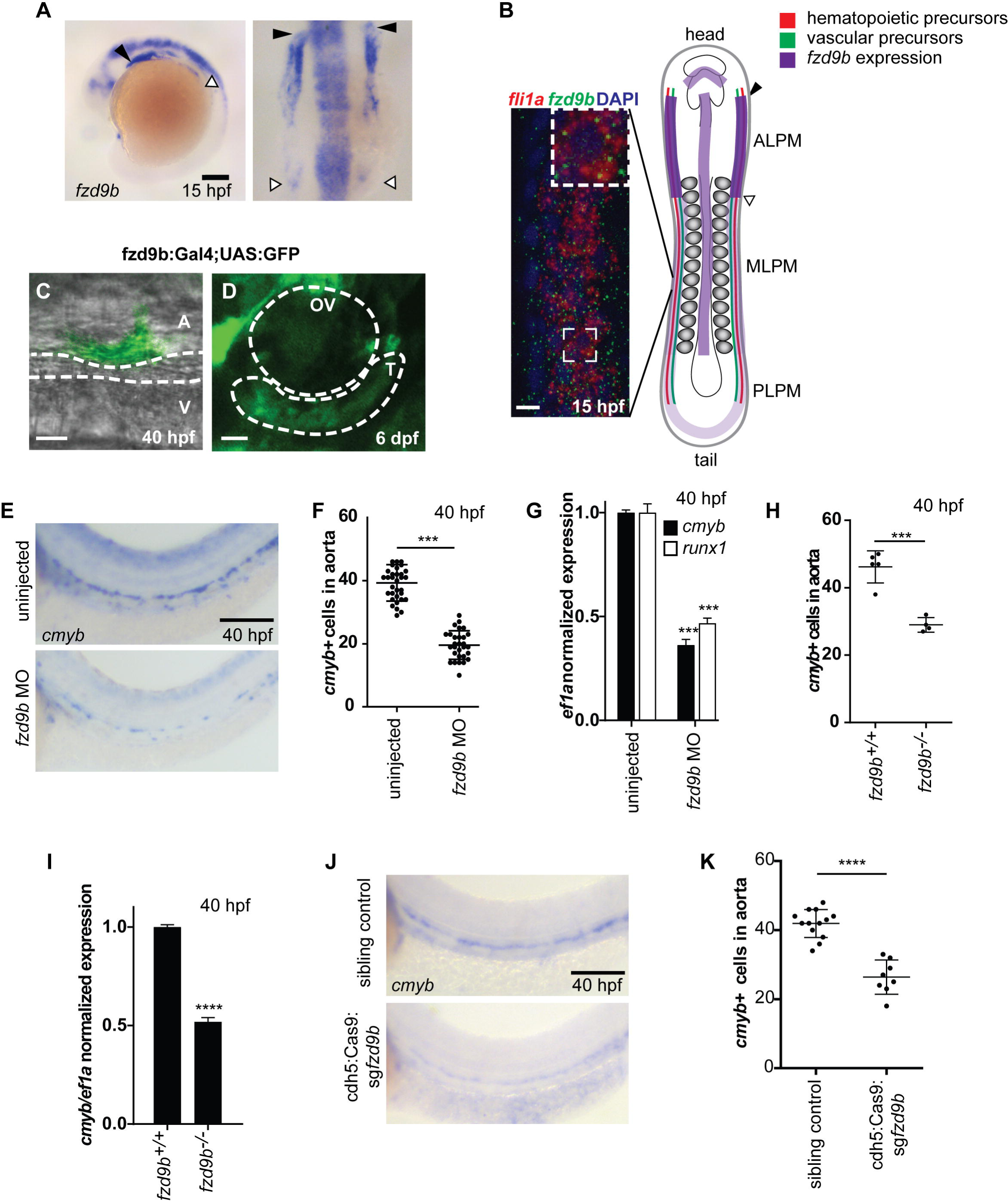
Fzd9b is required for zebrafish hematopoietic stem cell development. **A.** *fzd9b* is expressed in the anterior lateral plate mesoderm and neural tissues. Scale bar is 200um. **B.** Expression of *fzd9b* in 16 hpf zebrafish embryos overlaps with *fli1a* expression in the medial lateral plate mesoderm (MLPM), the tissue from which HSPCs are derived. PLPM-posterior lateral plate mesoderm, ALPM-anterior lateral plate mesoderm. Scale bar is 10um. **C.** Endothelial cell emerging from the dorsal aorta is labeled by *fzd9b:Gal4; UAS:GFP* at 40 hpf, during the time of HSPC emergence. Scale bar is 10um. **D.** Cells in the thymus are labeled by *fzd9b:Gal4; UAS:GFP* at 6 dpf. T-thymus, OV-otic vesicle; scale bar is 25um. **E.** WISH for the later HSC marker *cmyb* at 40 hpf in *fzd9b* morphants and controls. Scale bar is 30um. **F.** Quantification of **E**. **G.** qPCR for the later HSC marker *cmyb* in *fzd9b* morphants and controls at 40 hpf. Scale bar is 30um. **H.** Quantification of WISH for the later HSC marker *cmyb* at 40 hpf in *fzd9b* mutants and controls. **I**. qPCR for the HSC marker *cmyb* in *fzd9b* mutants and controls at 40 hpf. n.s. not significant. ***P<0.001, ****P<0.0001 compared to control (uninjected or WT siblings). (See also Figure S2).

The initial specification of HSPCs in *wnt9a* morphants is normal at 28 hpf, but later proliferative events are disrupted around 32-40 hpf (Grainger et al., 2016b). These phenotypes would be expected to be recapitulated by disruption of the Wnt9a receptor. To assess the function of Fzd9b in hematopoiesis, we used a translation-blocking morpholino (MO), which was able to block translation of a fluorescently labelled Fzd9b *in vivo* (Fig. S2C-D). Consistent with *wnt9a* loss of function, WISH for the early HSC marker *runx1* indicated that hematopoietic specification was not affected in *fzd9b* morphants (Fig. S2E). We confirmed these findings using reverse transcription-quantitative PCR (qPCR) on trunks and tails of morphants and sibling controls (Fig. S2F), indicating that Fzd9b is not required for hematopoietic specification. The expansion of hematopoietic cells is defective in *wnt9a* mutants and morphants, as determined by WISH for HSPC markers at 40 hpf (Grainger et al., 2016b); likewise, *cmyb* positive cells were significantly reduced in the dorsal aorta of *fzd9b* morphants at 40 hpf. These findings were consistent with a reduction of *cmyb* and *runx1* at 40 hpf by qPCR (Fig. 2E-G). In addition, loss of *fzd9b* led to a loss of emerging HSPCs, as indicated by confocal imaging of *gata2b/kdrl* double positive cells in the floor of the dorsal aorta at 46 hpf (Fig. S2G-I) (Butko et al., 2015). This loss of early HSPCs persisted throughout development, where *rag1*+ thymocytes (T cell precursors derived from HSPCs) were reduced in *fzd9b* morphants (Fig. S2J-K). This effect was specific to HSPCs, as markers for aorta (*dlc, dll4, notch1b*), vasculature (*kdrl*) and pronephros (*cdh17*) were normal (Fig. S2L-U). The *fzd9b* MO could be rescued with *fzd9b* mRNA (Fig. S2V-Y), suggesting that the effects of the MO were specific.

To validate our MO findings, we used CRISPR/Cas9 to generate two germline mutants of *fzd9b*, each of which had predicted premature stop codons early in the *fzd9b* coding sequence, harboring either a 2-base insertion, or a 7-base deletion, and both predicted to produce severely truncated N-terminal proteins of approximately 30 amino acid residues. Consistent with our MO findings, *fzd9b* mutants had a reduction in *cmyb*+ cells at 40 hpf (Fig 2H-I). Finally, we used a previously described transgenic approach where a guide RNA to *fzd9b* is expressed ubiquitously and *Cas9* expression is spatially regulated by *UAS* to conditionally inactivate *fzd9b* in early endothelial cells (Ablain and Zon, 2016), which also led to a reduction in the number of *cmyb*+ cells at 40 hpf (Fig. 2J-K), indicating that Fzd9b is required in the endothelium for HSPC development. Taken together, these results indicate Fzd9b is required for HSPC development, downstream of fate specification, and specifically in endothelial cells.

### Fzd9b interacts genetically with Wnt9a

We previously showed that Wnt9a’s effect on hematopoietic development required β-catenin (commonly referred to as the canonical Wnt pathway)(Grainger et al., 2016b). We therefore hypothesized that loss of *fzd9b*, the putative Wnt9a receptor, could be rescued with overexpression of constitutively-active (CA) β-catenin. Indeed, expression of CA-β-catenin under regulatory control of a *gata2b* promoter region (expressed exclusively within HE) was sufficient to rescue loss of *fzd9b* (Fig. 3A-E), indicating that like Wnt9a, Fzd9b functions upstream of β-catenin. The assembly of Fzd-Lrp5/6 heterodimers in response to a Wnt ligand is critical to enable signal transduction in β-catenin dependent signaling (Tamai et al., 2000). Consistent with Wnt9a-Fzd9b operating upstream of β-catenin, the Wnt9a-Fzd9b signal could be synergistically increased *in vitro* in cells co-transfected with Lrp6 (Fig. S3A). To determine if Wnt9a-Fzd9b signaling was reliant on endogenous LRP6 expression, we generated a HEK293T STF line deficient for LRP6 using CRISPR/Cas9. The resultant cell line harbored three alleles with large deletions in exon 2 of LRP6, which encodes part of the first extracellular beta-propeller, leading to a complete loss of protein as assessed by immune-blotting (Fig. 3F). These LRP6-deficient cells were compromised in their ability to activate STF reporter activity upon Wnt3a addition (Fig. S3B); however, treatment with GSK3 inhibitor (CHIR98014), which activates downstream signaling independent of Wnt-Fzd-LRP interactions, stimulated STF activity, indicating that downstream signaling components were intact (Fig. S3C). Consistent with these results, Wnt9a-Fzd9b signaling also required LRP6 (Fig. 3G). Therefore, the Wnt9a-Fzd9b complex operates upstream of β-catenin.

**Figure 3:**
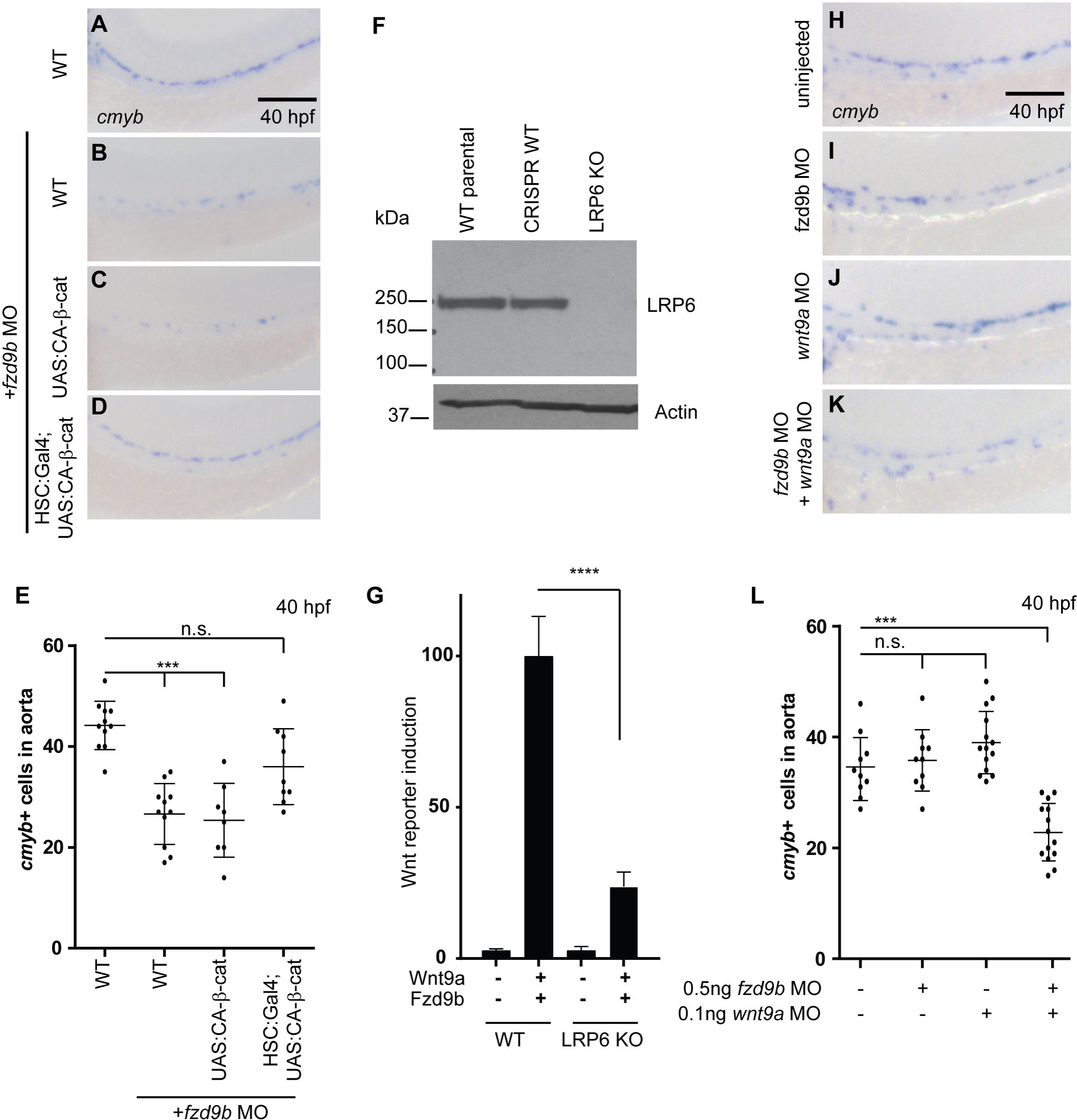
Fzd9b interacts genetically with Wnt9a. **A-D.** WISH for the HSC marker *cmyb* at 40 hpf in WT (**A, B**), *UAS:CA-β-catenin* (**C**) and *gata2b:Gal4;UAS:CA-β-catenin* (**D**), injected with *fzd9b* morpholino (**B-D**). Scale bar is 30um. **E.** Quantification of **A-D**. **F.** LRP6 immunoblot of lysates from WT HEK293T STF cells (WT parental), CRISPR-treated cells without disruption of LRP6 (CRISPR WT), and LRP6 null mutant line (LRP6 KO). **G.** Wnt reporter induction in WT and LRP6 KO HEK293T STF cells transfected with Wnt9a and Fzd9b. **H-K.** WISH for the HSC marker *cmyb* at 40 hpf in *wnt9a* (**J, K**) and *fzd9b* (**I, K**) low dose MO injected fish and sibling controls (**H**). Scale bar is 30um. **L.** Quantification of **H-K**. n.s. not significant. ***P<0.001 compared to WT controls. (See also Figure S3).

To confirm that Wnt9a and Fzd9b function in the same pathway *in vivo*, we used genetic non-complementation with suboptimal MO dosages. We found that a low dose of either Wnt9a or Fzd9b MO was not sufficient to affect hematopoiesis, while compound morphant animals had a reduction in *cmyb*+ cells similar to either *wnt9a* or *fzd9b* loss of function, providing genetic evidence that these components operate in the same genetic pathway (Fig. 3H-L). Altogether, these data indicate that Wnt9a and Fzd9b function in the same genetic pathway, upstream of β-catenin.

### WNT9A and FZD9 contribute to human hematopoietic development *in vitro*

Having identified a specific Wnt9a-Fzd9b signal *in vivo* in zebrafish, we sought to identify an interaction partner for human WNT9A. Using the STF reporter assay as above, we determined that only FZD9 coupled effectively with WNT9A (Fig. 4A), indicating that this signaling specificity is conserved from fish to human. To determine the requirement for WNT9A and FZD9 in human hematopoietic development, we employed an established differentiation protocol to drive human embryonic stem (hES) cell lines towards hematopoietic fates (Ng et al., 2008) (Fig. 4B). Previously, we showed that WNT9A promotes this differentiation, as monitored by expression of CD34, CD45 and RUNX1 (Richter et al. 2018). Disrupting WNT9A or FZD9 expression using short hairpin RNAs (shRNAs) significantly compromised the ability of hES cells to generate CD34/CD45 double positive HSPCs (Fig. 4C-F), suggesting that WNT9A and FZD9 contribute to human HSPC derivation. These results were confirmed with qPCR for endothelial (*CD34*) and hematopoietic markers (*CD31*, *CD45*, *CMYB*) (Fig. S4A-D). Differences in the abilities of these cells to differentiate were not due to loss of pluripotency of the undifferentiated hES cells, as they still abundantly expressed the pluripotency markers TRA1-81 and SSEA4 (Cooper et al., 2002; Thomson et al., 1998)(Fig. S4E-G). Thus, WNT9A and FZD9 are required for *in vitro* HSPC development in human cells.

**Figure 4:**
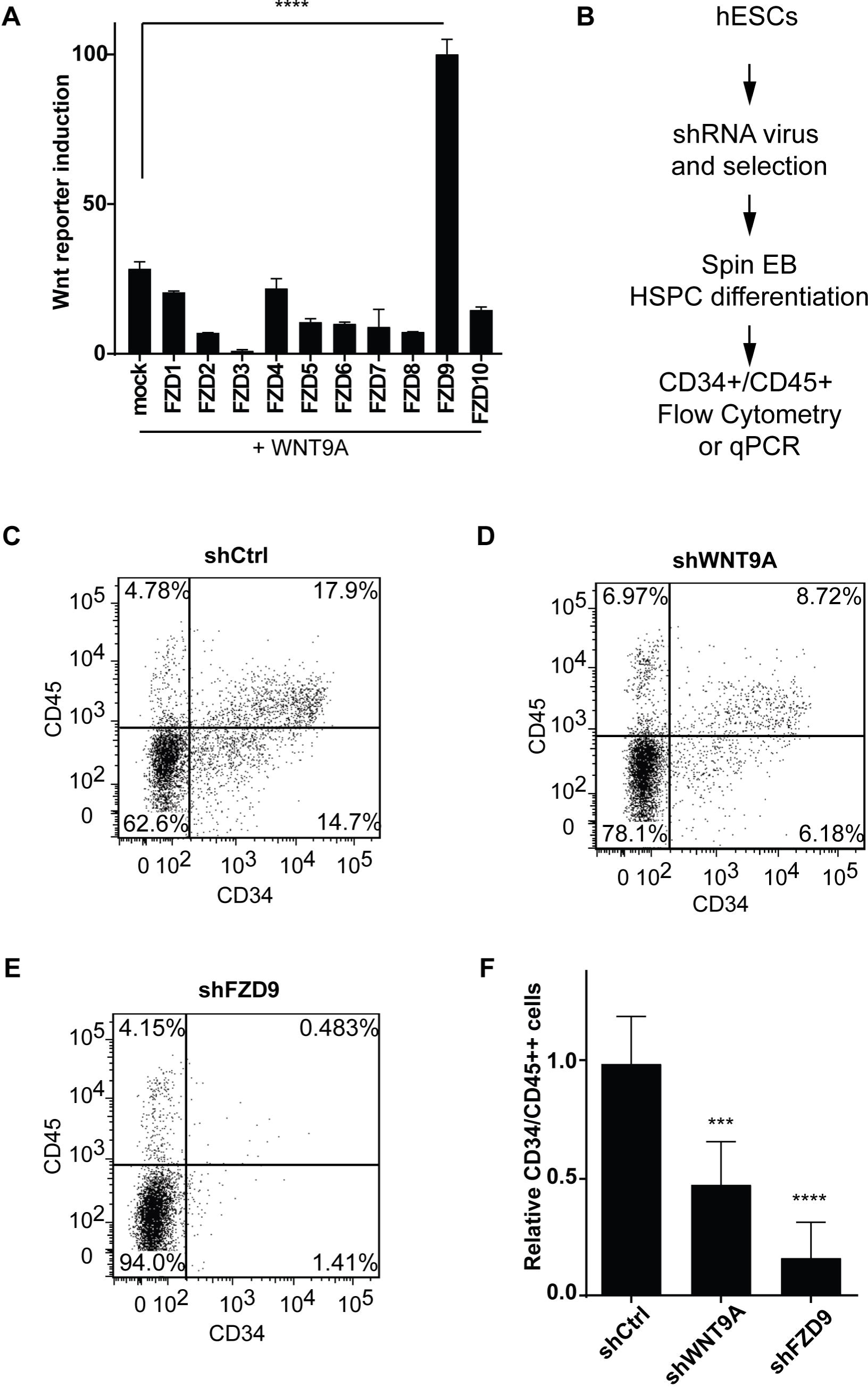
WNT9A and FZD9 are required for human hematopoiesis *in vitro*. **A.** Quantification of HEK293T cell STF assay screen with human WNT9A and FZDs. Bars represent the mean of a minimum of 3 biological replicates; error bars represent standard deviation; *P<0.05, **P>0.001, ****P<0.0001 compared to WNT9A alone (mock). **B.** Schematic of experimental design for HSPC derivation. Representative flow cytometry plots of CD34 vs CD45 cells after 14 days of differentiation towards HSPCs in shControl (**C**), shWNT9A (**D**) and shFZD9 **(E)** transduced cells. Note the loss of double positive cells with the loss of WNT9A or FZD9. **F.** Quantification CD34/CD45 double positive cells. Bars represent the mean from a minimum of 6 biological replicates from 2 independent experiments from **C-E**. *P<0.05, ***P<0.001 compared to shControl cells. (See also Figure S4).

### Wnt9a-Fzd9b specificity is mediated intracellularly

How the Wnt-Fzd signaling complex relays its signal and establishes specificity is poorly understood. One possibility is that Wnts engage the extracellular domains of Fzds with varying affinities. The crystal structure of a Wnt in complex with the extracellular domain of Fzd indicated that the Wnt molecule makes multiple contacts with the cysteine rich domain (CRD) (Janda et al., 2012). Furthermore, a number of studies showed that Wnts interact with varying affinities with the CRD (Dijksterhuis et al., 2015; Rulifson et al., 2000). Using the STF assay system and zebrafish cDNAs, we assessed the requirement for the Fzd9b CRD in mediating the Wnt9a signal *in vitro* and found that in the absence of the Fzd9b CRD, the Wnt9a-Fzd9b signal was attenuated (Fig. 5A), consistent with previous findings that the CRD is required for Wnt9a-Fzd9b signaling.

**Figure 5:**
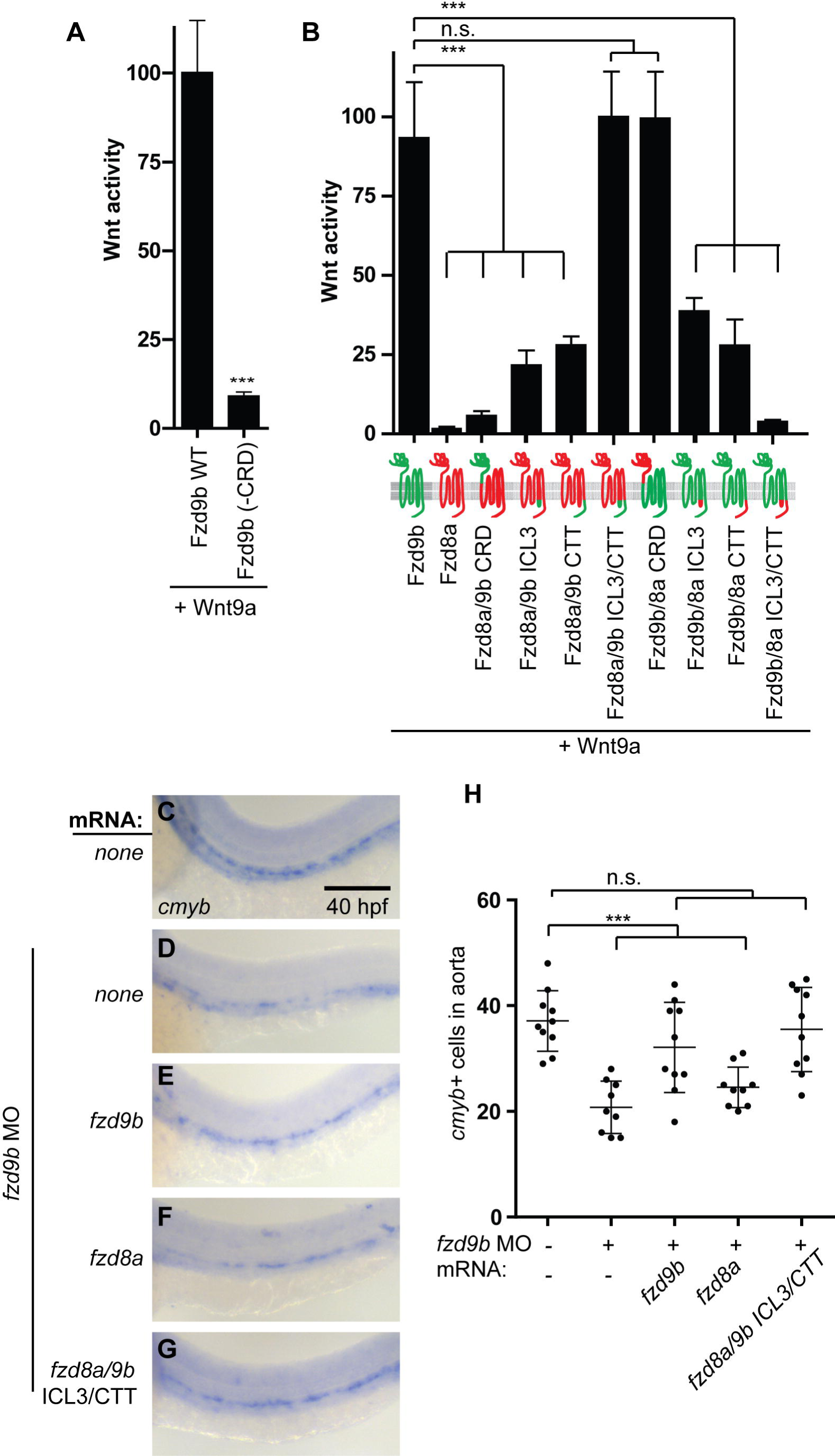
Wnt9a-Fzd9b specificity is mediated intracellularly. **A.** Quantification of HEK293T cell STF assay with zebrafish Wnt9a and Fzd9b with and without the cysteine rich domain (CRD). **B**. Quantification of HEK293T cell STF assay with zebrafish Wnt9a and Fzd9b/Fzd8a chimeras. Schematic representations of the constructs used are shown on the x-axis. Red represents coding regions from *fzd8a*; green represents coding regions from *fzd9b*. Bars represent the mean of a minimum of 3 biological replicates; error bars represent standard deviation; ***P<0.001 compared to Wnt9a with WT Fzd9b. **C-F.** WISH for the HSC marker *cmyb* at 40 hpf in *fzd9b* morphants (**D-G**) injected with mRNAs for *fzd9b* (**E**), *fzd8a* (**F**) and *fzd8a* with ICL3 and CTT from *fzd9b* (**G**) and uninjected control (**D**). Scale bar is 30um. **H.** Quantification of **D-G**. n.s. = not significant. ***P<0.001 compared to WT controls. (See also Figure S5).

We next sought to determine which domain(s) are required for Wnt9a-Fzd9b signaling specificity by constructing a series of chimeric Fzd transgenes between Fzd9b and Fzd8a, a Fzd that did not couple with Wnt9a to activate STF reporter activity (Fig. 1A). Surprisingly, a chimeric receptor in which the CRD of Fzd8a was replaced with that of Fzd9b did not signal (Fig. 5B), while the chimeric Fzd9b receptor carrying the Fzd8a CRD produced wild-type Wnt9a signaling activity (Fig. 5B), suggesting that the identity of the CRD is not a critical determinant in coupling Wnt9a-Fzd9b signaling. Furthermore, these results suggest that other domains of Fzd are critical in conferring Wnt9a specificity.

The signaling events downstream of Wnt-Fzd interaction and oligomerization with Lrp are poorly understood but are thought to involve interaction with intracellular mediator proteins such as Disheveled (Dsh), which is thought to interact with Fzd at the third intracellular loop (ICL3) and the C-terminal tail (CTT) (Tauriello et al., 2012; Wong et al., 2003). To assess the role of the ICL3 and CTT in mediating Wnt9a-Fzd9b signaling, we generated further Fzd9b-Fzd8a chimeric constructs (Fig. 5B). Using this approach, we found that substituting out either the ICL3 or CTT from Fzd9b with those of Fzd8a was sufficient to dampen signaling, while introducing these domains from Fzd9b into Fzd8a modestly increased the signal (Fig. 5B). Substituting both the ICL3 and CTT from Fzd9b with those from Fzd8a completely ablated the signal, and conversely substituting these Fzd9b domains into Fzd8a was sufficient to produce wild-type signaling levels (Fig. 5B), suggesting that signaling specificity for Wnt9a-Fzd9b lies entirely within the ICL3 and CTT domains. We were also able to recapitulate these findings using zebrafish and human cDNAs encoding chimeras for Fzd9b/FZD9 and Fzd4/FZD4 in our STF assay (Fig. S5A-B), indicating that the ICL3 and CTT are required for zebrafish Fzd9b and human FZD9 signaling.

As the Fzd protein matures and is exported to the cell surface, post-translational modifications including glycosylation increase the molecular weight (Janda et al., 2012; MacDonald and He, 2012; Yamamoto et al., 2005). Using V5-tagged Fzd constructs, we found that there was no change in the ability of these chimeric constructs to mature or be modified post-translationally (Fig. S5C). To confirm that the chimeric Fzd9b constructs that did not signal were transported to the cell surface, we performed immunofluorescence (IF) with a Fzd9b antibody directed to the extracellular the region between the CRD and the first transmembrane domain of Fzd9b (Fig. S5D). IF using this antibody on non-permeabilized cells confirmed that chimeric Fzd9b proteins were expressed on the cell surface at levels similar to wild-type Fzd9b (Fig. S5E). We further confirmed these findings using flow cytometry (Fig. S5F), indicating that differences in signaling were not due to differences in Fzd protein expression, maturation, or transport to the cell surface.

Altogether, these data indicated that ICL3 and CTT mediate the Wnt9a-Fzd9b signaling specificity *in vitro*. To determine if these domains were able to fulfill Fzd9b function in HSPC development, we co-injected mRNAs for *fzd9b, fzd8a* or *fzd8a* with the ICL3 and CTT from *fzd9b* in the context of *fzd9b* MO and found that only *fzd9b* and *fzd8a* with the ICL3 and CTT from *fzd9b* were able to rescue loss of *fzd9b* (Fig. 5C-H). Taken together, these data indicate that Wnt9a-Fzd9b specificity is regulated by the intracellular ICL3 and CTT domains of Fzd9b.

### Wnt9a, Fzd9b and EGFR form a complex

Since the above data indicate that Wnt9a-Fzd9b specificity is mediated intracellularly, we postulated the existence of another signaling component that spanned the membrane and contacted both Wnt9a and the intracellular portion of Fzd9b. Since we were able to analyze Wnt9a-Fzd9b signaling specificity using zebrafish cDNAs in human cells, we furthermore hypothesized that this signaling component would be a highly conserved. Therefore, to determine which proteins are recruited to the ICL3 and CTT of Fzd9b in response to Wnt9a, we generated a stable HEK293T cell line harboring Fzd9b fused to the peroxidase APEX2 (Lam et al., 2015). In the presence of hydrogen peroxide and biotin-phenol, endogenous proteins proximal to APEX2 (generally within 30 nm) are biotinylated, allowing for their enrichment with streptavidin beads, and subsequent identification by mass spectrometry (Lam et al., 2015). Due to the short labeling time required (60 seconds), it is possible to generate a timeline of proteins recruited to Fzd9b in response to Wnt9a. The APEX2 labelling from these cells was specific to biotin-phenol and hydrogen peroxide induction (Fig. S6A). Notably, the fusion protein did not interfere with membrane localization of Fzd9b (Fig. S6B-C) and was able to signal in response to Wnt9a as determined with STF reporter activity (Fig. S6D).

Gene ontology (GO) analysis of our APEX data revealed that in response to Wnt9a, the most changed biological processes included ERBB signaling (Fig. S6E). Consistent with this observation, the transmembrane proteins most enriched by proximity labeling were EGFR (also known as ERBB1), and ERBB2 (Fig. S6F). Disrupting EGFR expression in HEK293T cells stably expressing Fzd9b with short interfering RNA (siRNA) reduced cell surface binding of Wnt9a (Fig. 6A-H), suggesting that EGFR plays a role in the Wnt9a-Fzd9b interaction. Consistent with this observation, the EGFR ligand blocking antibody Cetuximab (Doody et al., 2007) dampened the Wnt9a-Fzd9b signal (Fig. 6I). These data are consistent with a model in which EGFR forms a complex with Wnt9a and Fzd9b to transmit the Wnt signal.

**Figure 6:**
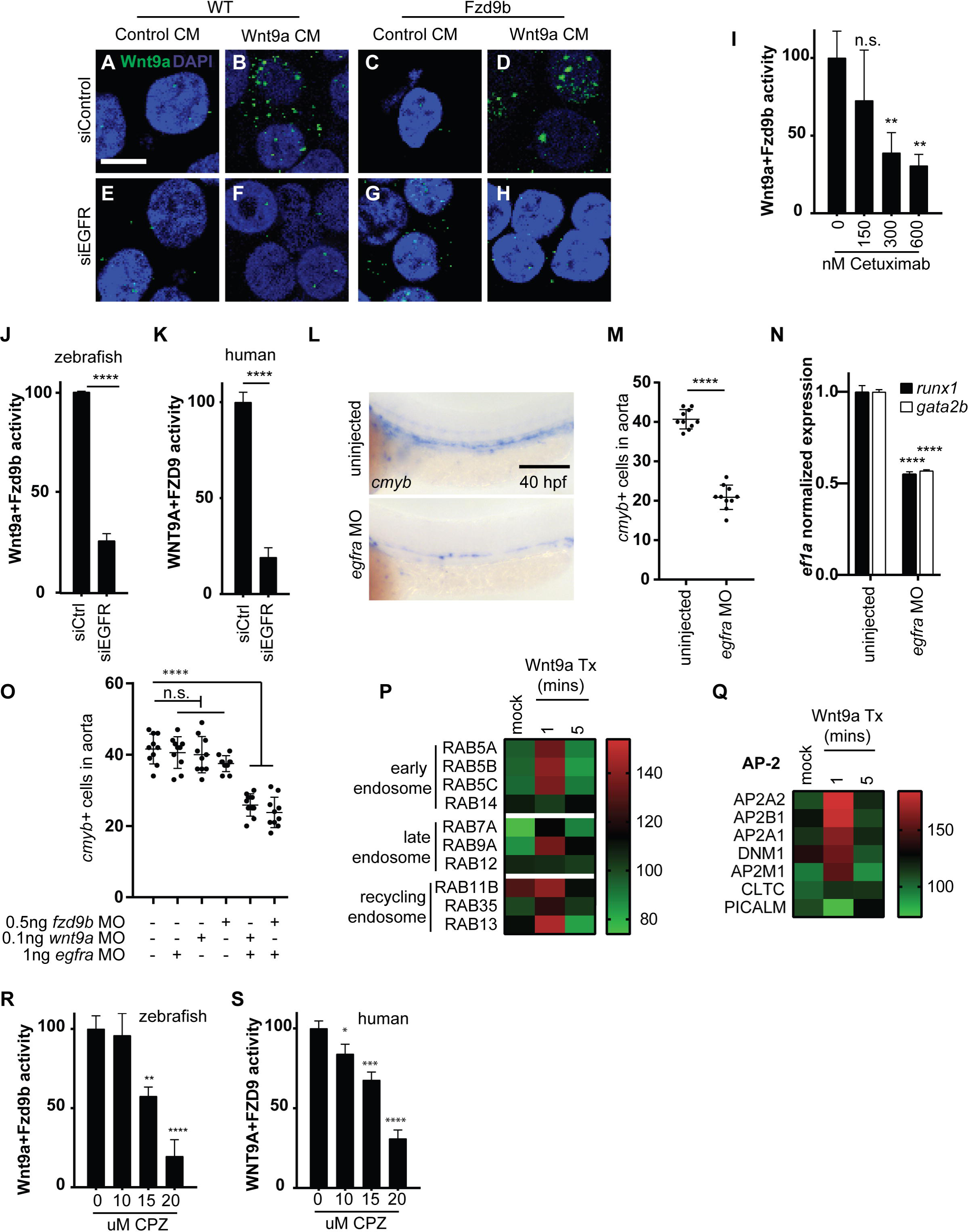
EGFR mediates Wnt9a-Fzd9b signaling. **A-H.** Non-permeabilized immunofluorescence for Wnt9a in Fzd9b cells treated as shown. Scale bar is 15 um. **I.** Quantification of HEK293T cell STF assay with zebrafish Wnt9a and Fzd9b treated with the EGFR blocking antibody Cetuximab. **J.** Quantification of HEK293T cell STF assay with zebrafish Wnt9a and Fzd9b treated with control or EGFR siRNAs. **K.** Quantification of STF assay with human WNT9A CHO cells co-cultured with stably transfected FZD9 HEK293T STF cells treated with control or EGFR siRNAs. **L.** WISH for the later HSC marker *cmyb* at 40 hpf in *egfra* morphants and controls. Scale bar is 30um. **M.** Quantification of **L. N.** qPCR for the HSC markers *runx1* and *gata2b* in *egfra* morphants and controls at 40 hpf. **O.** Quantification of WISH for *cmyb* at 40 hpf in fish injected with suboptimal MO dosages. **P.** Heatmap of Fzd9b-APEX2 proximity labeled normalized intensities of RAB members in the early endosome, late endosome and recycling endosome over time. **Q.** Heatmap of Fzd9b-APEX2 proximity labeled normalized intensities of AP-2 members, as well as associated machinery. Quantification of STF assay with zebrafish Wnt9a and Fzd9b (**R**) or human WNT9A and FZD9 (**S**), cultured with concentrations of chlorpromazine (CPZ) as shown. In all figures, bars represent the mean of a minimum of 3 biological replicates; error bars represent standard deviation; *P<0.05, **P<0.01, ****P<0.0001 compared to controls. (See also Figure S6).

### EGFR is required for the Wnt9a-Fzd9b signal

Genes from ERBB family, including EGFR, encode single-pass transmembrane receptor tyrosine kinases that homo- and hetero-dimerize in response to multiple ligands and to stimulate a number of signaling cascades (Mishra et al., 2017). HEK293T cells transfected with a short interfering RNA (siRNA) to EGFR compromised the ability of both zebrafish and human Wnt9a/WNT9A and Fzd9b/FZD9 to stimulate STF reporter activity (Fig. S6G, Fig 6J-K). Similarly, using a previously validated MO to *egfra* (Zhao and Lin, 2013), we observed a decrease in the number of HSPCs at 40 hpf, similar to the phenotypes of *fzd9b* or *wnt9a* loss of function, and consistent with a role for Egfr in regulating the Wnt9a-Fzd9b signal (Fig. 6L-N). In addition, suboptimal MO dosing indicated that both Fzd9b and Wnt9a operate in the same pathway as Egfr during HSPC development (Fig. 6O). Furthermore, treatment of cells with the selective EGFR tyrosine kinase inhibitor AG1478 (Goishi et al., 2003; Osherov and Levitzki, 1994) significantly attenuated STF reporter activity of both zebrafish Wnt9a/Fzd9b and human WNT9A/FZD9 (Fig. S6H-I). Consistent with these results, EGFR kinase activity was also required for HSC development in zebrafish, as assayed by expression of the HSC markers *runx1* and *gata2b* at 40 hpf (Fig. S6J). These data indicate EGFR and its kinase activity are required for the Wnt9a-Fzd9b signal.

EGFR is known to cross-talk with other seven-pass transmembrane receptors such as G-protein coupled receptors (GPCRs), the consequences of which have effects on signal transduction, as well as receptor internalization and trafficking (Daub et al., 1997; Gschwind et al., 2001; Kue et al., 2002; Prenzel et al., 1999; Schafer et al., 2004; Tomlins et al., 2005; Vacca et al., 2000). Consistent with this, GO analysis of our APEX data indicated that the most enriched cellular component was “clathrin-coated endocytic vesicle” (Fig S6K). The APEX data also showed enrichment for proteins associated with early endosomes (RAB5A, RAB5B, RAB5C and RAB14), late endosomes (RAB7A, RAB9A and RAB12), and recycling endosomes (RAB11B, RAB35 and RAB13) (Fig. 6P), consistent with Fzd9b internalization in response to Wnt9a (Fig. 6I). Internalization of seven-pass transmembrane proteins can be mediated by clathrin- or caveolin-mediated endocytosis, as can Fzd-Wnt complexes (Blitzer and Nusse, 2006; Yamamoto et al., 2006). We observed the recruitment of members of the AP-2 complex and clathrin-mediated endocytosis machinery to Fzd9b in response to Wnt9a, indicating that internalization was mediated by clathrin (Fig. 6Q). Finally, both the zebrafish Wnt9a/Fzd9b and human WNT9A/FZD signals required clathrin-mediate endocytosis, as indicated by STF assay after treatment with the clathrin inhibitor chlorpromazine (Fig. 6R-S). These results indicate that Fzd9b is internalized and sorted through the endosome-lysosome in response to Wnt9a.

The C-terminal tail of Fzd9b contains two tyrosine (Y) residues, at 556 and 571 (Fig. 7A), which are predicted to be potential kinase substrates (Blom et al., 1999). Additionally, the Y556 on Fzd9b is highly conserved among vertebrates and Y571 is partially conserved, indicative of putative functional importance (Fig. S7A). Consistent with these predictions, treatment with Wnt9a increased Y-phosphorylation of Fzd9b (Fig. 7B). This increase was not observed in the presence of the EGFR tyrosine kinase inhibitor AG1478 (Fig. S7D), indicating that EGFR kinase activity is required to increase Y-phosphorylation on Fzd9b in response to Wnt9a. In addition, mutation of the Y556 (but not Y571) putative phosphorylation sites decreased the *in vitro* signaling capacity of Wnt9a (Fig. 7C). Finally, mutation at the corresponding tyrosine in human FZD9, Y562F, also led to a decrease in its signaling capacity (Fig. S7D), suggesting that this function is conserved. Together, these data indicate that Fzd9b is phosphorylated on tyrosine residue 556 in response to Wnt9a, which is required for its downstream signal.

**Figure 7:**
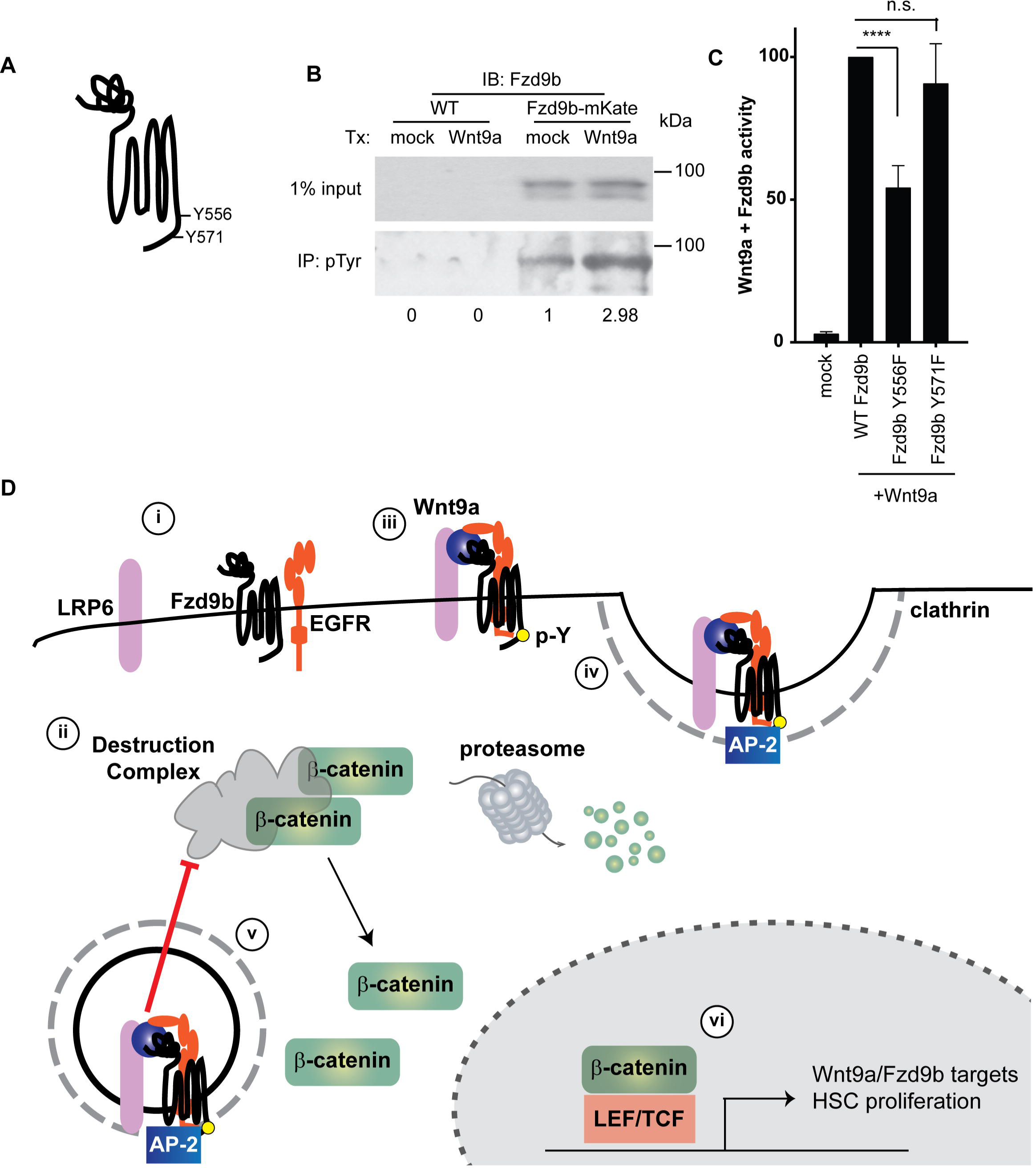
Fzd9b Tyrosine phosphorylation is required for Wnt9a-Fzd9b signaling. **A.** Schematic of Fzd9b protein with putative EGFR tyrosine phosphorylation sites.**B.** Western blot for Fzd9b in phosphotyrosine immunoprecipitations from WT or HEK293T cells stably transfected with Fzd9b-mKate and treated for 1 minute with either WT CHO conditioned medium, or Wnt9a-expressing CHO conditioned medium as shown. Note the increase in phosphotyrosine Fzd9b (2.98 vs 1, arbitrary units). **C.** Quantification of HEK293T cell STF assay with zebrafish Fzd9b point mutants. Bars represent the mean of a minimum of 3 biological replicates; error bars represent standard deviation; ****P<0.0001 compared to controls. **D.** Model of Fzd9b signaling specificity with Wnt9a. **i.** Fzd9b, LRP6 and EGFR are resident in proximity at the cell surface; **ii.** the destruction complex targets β-catenin for degradation in the absence of a ligand; **iii.** in the presence of Wnt9a, EGFR phosphorylates the Fzd9b tail at Y556 and Y571. **iv.** AP-2 and clathrin are recruited. **v.** Fzd9b-LRP6 oligomerization leads to dissociation of the destruction complex and release of β-catenin into the cytosol, allowing for nuclear translocation. **vi.** Nuclear β-catenin transactivates a program for HSC proliferation. (See also Figure S7).

Taken together, our data suggest a mechanism for the specificity of Wnt9a-Fzd9b signaling. Fzd9b, LRP6 and EGFR are resident in proximity at the cell surface (Fig. 7Di); in the absence of a ligand, β-catenin is targeted for proteasomal degradation by the destruction complex (Fig. 7Dii); in the presence of Wnt9a, these are bridged, allowing EGFR-mediated phosphorylation of the Fzd9b tail at Y556 (Fig. 7Fiii), leading to the recruitment of AP-2 and clathrin (Fig. 7Div). Once inside the cell, Fzd9b-LRP6 oligomerization leads to dissociation of the destruction complex and release of β-catenin (Fig. 7Dv). Finally, nuclear β-catenin transactivates a program for HSC proliferation (Fig. 7Dvi).

## Discussion

### EGFR and Fzd9b are required for the Wnt9a signal in vitro and in vivo

We have found that Fzd9b is required to relay the Wnt9a signal during HSPC development in zebrafish; this finding shows conservation to human cells as well, where WNT9A and FZD9 are required to generate HSPCs from human pluripotent precursors. We have determined that the specificity of the Wnt9a-Fzd9b (and the WNT9A-FZD9) signal relies on intracellular domains, not the extracellular CRD, suggesting the existence of a transmembrane co-factor. With a proximity-labeling based mass spectrometry approach, we identified that EGFR and Fzd9b are closely associated in the cell; EGFR (and specifically the tyrosine kinase activity) is also required for HSPC development and for the Wnt-Fzd9b signal. Altogether, these results suggest a general paradigm where particular Wnt-Fzd interactions mediate specific signals that are transduced in the context of particular co-receptors, such as receptor tyrosine kinases.

Our previously characterized requirement for Wnt9a in HSPC development had established that Wnt9a plays a specific role in the early amplification of HSPCs, which could not be rescued by other Wnts (Grainger et al., 2016b). Here we showed that this extracellular Wnt9a signal is relayed *in vitro* and *in vivo* by Fzd9b (or FZD9 in humans) and not by any other Fzd receptor. The requirement for Fzd9 in blood development is in line with previous data from mouse models, where loss of Fzd9 leads to a partially cell-intrinsic hematopoietic defect (Ranheim et al., 2005). Although the authors did not examine HSPCs directly, they did note several pathologies linked to HSPCs, such as depletion of B cells (one of the lymphoid lineages downstream of HSCs), splenomegaly, thymic atrophy and lymphadenopathy with age (Ranheim et al., 2005). These observations are consistent with a loss of HSPCs during development. Furthermore, disruption of *FZD9* expression has been linked to hematological malignancies, highlighting its importance in human hematopoietic cells as well (Jiang et al., 2009; Martin-Subero et al., 2009; Zhang et al., 2016). For instance, *FZD9* was identified as one of 6 genes most frequently hypermethylated in hematological neoplasms (Martin-Subero et al., 2009). In another example, *FZD9* was identified as the most frequently aberrantly methylated gene in a survey of 184 patients with myelodysplastic syndrome and acute myeloid leukemia; those with loss of heterozygosity of the *FZD9* allele had the poorest outcome (Jiang et al., 2009). Altogether, these data point to a role for Fzd9 in hematopoietic stem cell development.

The identification of EGFR contributing to HSPC development is surprising. There is little evidence for the involvement of EGFR in the hematopoietic niche. Loss of EGFR results in mid-gestational mortality in mice, precluding the analysis of HSPCs (Sibilia and Wagner, 1995). EGFR has been shown to promote the recovery of HSPCs in response to radiation injury (Doan et al., 2013); conversely, inhibition of EGFR promotes the mobility of HSPCs (Ryan et al., 2010). These are in general support of a role of EGFR in HSPCs that has not yet been explored, and consistent with our findings that it is required as a co-factor during HSPC development.

### A model for Wnt-Fzd signaling specificity

One long outstanding puzzle in the Wnt field has centered around the requirement for genetically encoding such a diverse set of ligands and receptors if Wnt ligands are as promiscuous as reported (Dijksterhuis et al., 2015; Ring et al., 2014; Yu et al., 2012). In addition, biochemical analysis of Wnt/Fzd interactions has been limited since Wnt purification is difficult, due to post-translational modifications making them highly hydrophobic (Willert et al., 2003). Finally, putative functional overlap between family members has made determination of specific function and cognate pairings hard to deconvolute using genetics (Ikeya et al., 1997; Ikeya and Takada, 1998; Yamaguchi et al., 2005). Therefore, potential pairings have remained elusive.

It has been hypothesized that Wnt-Fzd interactions are regulated by physical binding affinities. *Xenopus* Wnt8 interacts directly with the CRD of mouse Fzd8, as shown by x-ray crystallography (Janda et al., 2012). In addition, dissociation constant measurements of Wnt-CRD interactions revealed that different Wnt-Fzd combinations have different affinities and that many Wnts can physically interact with multiple (if not all) Fzd CRDs (Dijksterhuis et al., 2015; Hsieh et al., 1999; Ring et al., 2014; Yu et al., 2012). This has led to the prevailing model that Wnt ligands are promiscuous, signaling with any Fzd in proximity. This model has been supported in large part due to the fact that the prototypical ligand, Wnt3a was first to be purified, and is in fact able to activate a multitude of Fzd receptors *in vitro* (Dijksterhuis et al., 2015; Voloshanenko et al., 2017; Willert et al., 2003). In addition, the low solubility of Wnts leads to a very short secretion range, which predicts that Wnt-Fzd specificity is regulated largely at the level of spatio-temporal expression (Farin et al., 2016). Our results indicate that signaling specificity is regulated at an additional level involving the activation of co-receptors, though this does not preclude specificity contributions from secretion range, spatio-temporal localization or Wnt-Fzd affinities.

We have shown that Fzd9b is present in the membrane in proximity to its co-receptor EGFR, which has previously been shown to be membrane localized in an inactive conformation in the absence of a ligand (Purba et al., 2017). Once Wnt9a is present, EGFR phosphorylates tyrosine residues on the CTT of Fzd9b and leads to the recruitment of internalization machinery such as the AP-2 complex and clathrin. LRP6 is also required for this signaling event, as previously established for other β-catenin dependent signaling events (Baig-Lewis et al., 2007; Cong et al., 2004; Pinson et al., 2000; Schweizer and Varmus, 2003; Tamai et al., 2000; Wehrli et al., 2000). We therefore propose that the signaling specificity of a Wnt-Fzd pair is regulated (1) by the ability of Wnt to bind to the Fzd CRD, and (2) the recruitment of an activating co-receptor required for internalization (Fig. 7D). This combination of internalization and LRP recruitment then leads to nuclear β-catenin accumulation and target gene activation.

This model is in general agreement with the finding that Fzds have putative phosphorylation sites, and can be phosphorylated upon activation, though this has only been shown in the context of β-catenin independent signaling (Djiane et al., 2005; Shafer et al., 2011; Yanfeng et al., 2006). This is also in general agreement with the so called “endosomal Wnt signaling” model, which posits that Fzd-LRP6 oligomerization, phosphorylation and subsequent caveolin-mediated endocytosis are required for signaling, while clathrin-mediated endocytosis would result in signal dampening (Yamamoto et al., 2006; Yamamoto et al., 2008). In contrast, we have found that clathrin-mediated endocytosis is initiated following Fzd9b phosphorylation by EGFR, indicating that clathrin-mediated endocytosis can also lead to context-dependent signal activation. Of note, caveolin dependent endocytosis is characterized by the presence of caveolin-1, which we did not detect in our APEX data (Parton and Simons, 2007). This finding may also explain the puzzle of how *Drosophila,* which lack caveolin, but do have clathrin, are able to produce a Wnt signal (MacDonald and He, 2012), as well as how *caveolin*^-/-^ mice have augmented Wnt signaling in certain niches (Li et al., 2005; Sotgia et al., 2005). On the other hand, clathrin-mediated endocytosis has been shown to be required for *Drosophila* Wg (Wnt homologue) internalization and signaling *in vitro* (Blitzer and Nusse, 2006), which is also consistent with our findings. These differences in clathrin- and caveolin-mediated endocytosis leading to up- or down-regulation of signaling may relate back to the particular Wnt/Fzd/co-receptor combination activating their recruitment, since particular phosphorylation signatures are known to affect sorting of the endosome (Villasenor et al., 2016).

This model is also in agreement with findings of EGFR transactivation with other seven-pass transmembrane receptors such as GPCRs (Kose, 2017). Others have hypothesized that Fzd receptors are a sub-class of GPCRs (Dijksterhuis et al., 2014); there is limited, but growing evidence for Fzd receptor coupling to G proteins, although this is debated in the field (Ahumada et al., 2002; Halleskog et al., 2012; Katanaev, 2010; Katanaev et al., 2005; Kilander et al., 2014a; Kilander et al., 2011a; Kilander et al., 2011b; Kilander et al., 2014b; Liu et al., 2001; Liu et al., 1999; Sheldahl et al., 1999; Slusarski et al., 1997). Regardless of whether or not Fzds are true GPCRs, this does not preclude them from functioning in a manner similar to GPCRs. GPCR desensitization is regulated by internalization of the receptor, a process that is dependent on post-translational modifications such as phosphorylation (Jean-Charles et al., 2016). In our study, we have found that Fzd9b is phosphorylated on tyrosine residues in response to Wnt9a, consistent with a role for the receptor tyrosine kinase EGFR in this process. Altogether, these indicate that in response to Wnt9a, Fzd9b is phosphorylated by EGFR, and internalized through clathrin-mediated endocytosis. There is precedent for this particular mechanism of internalization of receptors. For example, trafficking of the GPCR CXCR7 requires serine/threonine phosphorylations at the C-terminus, followed by deubiquitination, which is likely required to allow interaction with AP-2 (Canals et al., 2012). Wnt signaling also depends on clathrin-mediated endocytosis in a context-dependent manner (Saito-Diaz et al., 2018). Finally, Wnt7a and Wnt7b signaling depend on the membrane anchored glycoprotein Reck and the GPCR Gpr124 together with Fzd4 (Cho et al., 2017; Eubelen et al., 2018; Zhou and Nathans, 2014), while Norrin and Fzd4 signaling relies on Tspan12 (Lai et al., 2017), supporting the notion that specificity of Wnt-Fzd pairs relies on co-receptor complexes. In particular, low level affinities of Reck for Wnt7 seems to be important for regulating the availability for Wnt7 to bind its cognate Fzd receptor, through oligomerization with Gpr124, Dishevelled and Fzd (Eubelen et al., 2018).

There are many families of receptor tyrosine kinases, including EGFR/ERBB, FGF receptors, vascular endothelial growth factor receptors and Eph receptors, among others. Of these, many are known to play complementary roles with Wnt signaling during development and cancer. For example, Wnt10b and FGF3 cooperate in mammary tumorigenesis (Lee et al., 1995), as do FGF8 and Wnt1 (MacArthur et al., 1995). In planarians, Wnt and FGFR are both required for anterior-posterior patterning of the brain (Kobayashi et al., 2007) and chondrocyte differentiation (Buchtova et al., 2015). In addition to these, there are overlapping functions for Wnt and FGF in the developing pre-somitic mesoderm, with consequences in somitogenesis and axial elongation (reveiwed in Mallo, 2016). It is conceivable that activation of these receptors in the context of a Fzd may also play a cooperative role in activation of the Wnt signal.

### Wnt-Fzd co-receptors in regenerative medicine and cancer

One context where knowledge of Wnt-Fzd and co-factor specific interactions will be critical will be in regenerative medicine, such as in the development of protocols to derive different tissues *in vitro* from pluripotent precursor cells. Although there is increasing evidence for HSC derivation *in vitro* (Sturgeon et al., 2013; Wahlster and Daley, 2016), the field still struggles with deriving HSCs *in vitro* from pluripotent precursors, using xenograft-free conditions. This would allow for patient-specific cell therapies for diseases of the blood such as leukemias, lymphomas, anemias, and auto-immune disorders. In large part, the requirement for β-catenin dependent Wnt signaling in these protocols has been substituted for with the prototypical ligand, Wnt3a, or with GSK3 inhibitors, which potently activate Wnt signaling. Our data suggest that certain biological functions of the Wnt signal cannot be substituted for with other Wnt agonists; for example, the loss of Wnt9a during early HSPC proliferation in zebrafish cannot be rescued by Wnt3a (Grainger and Richter et al, 2016b). Additionally, the complement of Fzd receptors and putative co-receptors expressed in the correct spatio-temporal location will be essential to recapitulating endogenous developmental cues in the dish. The hematopoietic field has struggled for decades to make *bona fide* HSCs *in vitro* from either conversion of non-hematopoietic cells or directed differentiation from pluripotent precursors. Many efforts to reprogram non-hematopoietic human or mouse cells directly to HSCs resulted in HSPC-like cells that could not be sustained *in vivo* (Batta et al., 2014; Doulatov et al., 2013; Elcheva et al., 2014; Pereira et al., 2013; Pulecio et al., 2014; Sandler et al., 2014). However, long-term, engraftable, immunocompetent HSCs can now be derived by direct conversion of mouse endothelial cells, using a set of four transcription factors using mouse cells (Lis et al., 2017). A similar approach has also been used in human cells derived from pluripotent precursors; however, the therapeutic use of these is challenged by exogenous expression of transcription factors (Sugimura et al., 2017). One potential issue may be overlooking this early proliferative event, for example, along with the requirement for FZD9 and EGFR as co-receptors in this process, as this strategy uses the general GSK3β inhibitor CHIR99021 (Dege and Sturgeon, 2017; Ditadi and Sturgeon, 2016; Sugimura et al., 2017). Taking these specific requirements into consideration will be critical to the advancement of regenerative medicine.

Cross talk between WNT and EGFR has been long observed, especially in cancer cells, though the molecular mechanisms regulating this process have been poorly understood, but are thought to operate at multiple levels (Paul et al., 2013). For instance, EGFR activating mutations are associated with better prognosis when coupled with unmethylated Wnt antagonist promoter regions (Suzuki et al., 2007), suggesting that these Wnt antagonists may influence EGFR activity. EGF-mediated activation of EGFR leads to an increase in β-catenin transcriptional activity in cancer cells (Ji et al., 2009), as does expression of activated variants of EGFR (Del Vecchio et al., 2012). Wnt1 and Wnt5a are thought to induce *cyclinD1* expression through EGFR activation in mammary HC11 epithelial cells, which could be blocked using a ligand blocking antibody to EGFR (Civenni et al., 2003); an alternate explanation to these findings could be aberrant EGFR-mediated endocytosis coupled to the Fzd receptor that is mediating the Wnt signal. It has also been suggested that this occurs through direct Y-phosphorylation of β-catenin by various receptor tyrosine kinases (Krejci et al., 2012), though the cooperation between these receptor tyrosine kinases and Wnt transcriptional output could also be re-interpreted at the level of the membrane.

EGFR and Wnt have also been suggested to cross-talk during normal development and homeostasis. For instance, EGFR signaling, similar to Wnt, is required for re-establishing the proximal-distal axis during leg regeneration in crickets, where RNAi loss of function phenocopies the loss of Wnt (Nakamura et al., 2008). There is also evidence for Wnt/EGFR function in Drosophila spiracle development (Maurel-Zaffran et al., 2010), hemocytes (Zettervall et al., 2004), glucose homeostasis (Cardone et al., 2012) and epithelial regeneration (Georgopoulos et al., 2014), among others.

Finally, determining how individual WNTs and FZDs are coupled will have important therapeutic implications. Overactivation of the WNT pathway is associated with a number of human malignancies, including (but not limited to) cancers of lungs, ovaries, breast, skin, brain and intestinal tract (Clevers and Nusse, 2012). In some subsets of these cancers, different FZD receptors are overexpressed, leading to an amplification of the WNT signal (Chan and Lo, 2018; Ueno et al., 2013). Importantly, pan-WNT inhibitory therapies cause toxic side effects, due to the importance of WNT signaling in other tissues, such as the maintenance of the intestinal epithelium (Korinek et al., 1998; Schepers and Clevers, 2012; van Es et al., 2012). Furthermore, our observation that Cetuximab, which blocks ligand binding to EGFR, disrupts Wnt9a/Fzd9 signaling suggests potential alternative mechanisms of action for this chemotherapeutic agent. Being able to derive therapies that are specific to WNT-FZD pairs will allow for more specific targeting of these cancer cells.

## Experimental Procedures

### Cell culture and luciferase reporter assays

HEK293T cells with a stably integrated Super-TOP-Flash reporter (STF) (Fernandez et al., 2014) and Chinese hamster ovary (CHO) cells were grown in Dulbecco’s modified Eagle’s medium (DMEM) supplemented with 10% heat-inactivated fetal bovine serum (FBS) under standard conditions.

Cells were seeded into six-well plates and transfected using polyethyleneimine (PEI), 50ng of *renilla* reporter vector, 200ng of Wnt, Fzd or Lrp6 expression vector, with a total of 1 ug of DNA/well. For co-culture experiments, cells were passaged 24 hours after transfection and plated together for analysis. For siRNA experiments, a 6 well was treated with 10pmol of siRNA using RNAiMax transfection reagent (Invitrogen). All human WNT9A cell culture experiments except for the initial screen were performed by co-culturing stably expressing WNT9A CHO cells with stably expressing FZD9 HEK293 STF cells. All transfected cells were harvested 48 h post-transfection and all conditioned medium or co-cultured cells were harvested 24 h post-treatment; the lysates processed and analyzed using the Promega Dual Luciferase Assay System according to the manufacturer’s instructions. Each experiment was performed with biological triplicate samples and reproduced at least one time with a similar trend.

10 cm plates of 293T cells were transfected with 10ug of constructs encoding chimeric Fzd cDNAs with a C-terminal V5 tag. For immunofluorescence, cells were plated onto glass coverslips after 24 hours, and stained with our Fzd9b antibody, generated to a region between the CRD and the first transmembrane domain, under non-permeabilized conditions, and according to standard protocols. For flow cytometry, cells were harvested with Accutase, pelleted, resuspended in PBS with 1% BSA and 1mM EDTA, filtered through an 80um filter and sorted using a BD Fortessa flow cytometer. For western blots, cells were harvested 48 hours after transfection in TNT buffer (1% Triton X-100, 150mM NaCl, 50mM Tris, pH8.0), with protease inhibitors. Westerns were performed according to standard procedures, using antibodies for V5 (1:5,000, GeneTex) and β-actin (1:20,000, Abcam).

For immunoprecipitation, cells were washed 3 times in PBS and lysed in radio immunoprecipitation assay (RIPA) buffer (10mM Tris-HCl, pH8, 10mM EDTA, 0.5mM EGTA, 1%Tx-100, 0.1% deoxycholate, 0.1% SDS, 140mM NaCl), supplemented with protease and phosphatase inhibitor tablets (Pierce) and 20mM N-ethylmaleimide, for 30 minutes at 4°C with rocking. Resultant lysates were cleared of cell debris by centrifugation at 15,000g for 10 minutes at 4°C and quantified by Bradford Assay. A minimum of 200ug of protein was diluted into 400uL total volume with RIPA buffer; 2ug of antibody was added and incubated at room temperature for 30-60 minutes. Antibody-protein complexes were precipitated using Protein a Dynabeads (Invitrogen) for 30 minutes at room temperature, washed 3 times with 1mL of RIPA buffer and eluted using Laemmli buffer at room temperature. Precipitates were run on a Western blot for Fzd9b. Western blot intensities from pulldowns were quantified by densitometry using ImageJ, normalized to input controls.

EGFR inhibition was performed using a 5mM stock of AG1478 in 50:50 Ethanol: DMSO. The final concentration used was 2.5uM. Clathrin-mediated endocytosis was inhibited using chlorpromazine from a 1M stock in 50:50 Ethanol: DMSO. Final concentrations used are as indicated in the figure. The Human EGFR phosphorylation Array C1 (Raybiotech) was carried out according to the manufacturer’s recommendations using lysates from Fzd9b-mKate HEK293T cells treated with Wnt9a or mock conditioned medium for 1 minute.

### Wnt9a surface binding assay

Conditioned medium was collected from stably expressing Wnt9a or parental CHO cells and concentrated 10X using a 30kDa molecular weight cutoff column. HEK293T cells stably expressing Fzd9b were transfected with siControl or siEGFR and plated on 0.1% gelatin coated glass coverslips after 24 hours; after a further 24 hours, cells were treated with cold conditioned medium for 3 hours at 4°C, rinsed with PBS and fixed with 4% PFA at 4°C for 20 minutes and at room temperature for 10 minutes. Immunofluorescence was performed using standard non-permeabilizing methods with a rabbit polyclonal antibody generated to zebrafish Wnt9a.

### Generation of LRP6 knockout HEK293T STF line

A confluent 100mm plate of HEK293T STF cells were transfected with 3mg each of Cas9 (Jao et al., 2013) and two guide RNAs under regulatory control of a U6 promoter. The guide RNAs (GGGCTTGGAGGATGCAGCTG and GGATCTAAGGCAATAGCTCT) targeted the second exon of LRP6. Single cell clones were validated for loss of LRP6 by sequencing the genomic locus, Western blot using a rabbit monoclonal antibody (1:1000, C47E12, Cell Signaling) and for loss of LRP6 function with STF assay for mouse Wnt3a, which is reliant on LRP6 for signaling (Tamai et al., 2000).

### Fzd9 and Wnt9a antibody generation

GST fusion proteins for immunogens of Fzd9b (provide amino acid numbers) and Wnt9a (provide amino acid numbers) were purified by standard methods. Rabbits were immunized with GST proteins, boosted and bled for serum according to standard methods (Lampire Biologicals). Antibodies for Wnt9a or Fzd9b were affinity purified against the same antigens fused to maltose binding protein (MBP), according to the manufacturer’s recommendations (Fisher), and stored in PBS/50% glycerol at −80°C.

### Zebrafish lines and maintenance

Zebrafish were maintained and propagated according to University of California and local Institutional Animal Care and Use Committee policies. AB*, *Tg(kdrl:Cherry-CAAX)^y171^, Tg(fli1a:eGFP)^zf544^, Tg(cdh5:Gal4)^mu101^, Tg(UAS:CA-β-catenin)^sd47Tg^ Tg(gata2b:KalTA4^sd32^; UAS:Lifeact:eGFP^mu271^), wnt9a^∆28/∆28,sd49^, Tg(UAS:Cre)^t32240Tg^ Tg(bactin2:loxP-BFP-loxP-DsRed)^sd27Tg^* lines have been previously described (Bertrand et al., 2010; Bertrand et al., 2008; Bouldin and Kimelman, 2014; Butko et al., 2015; Espin-Palazon et al., 2014; Grainger et al., 2016b; Jin et al., 2005; Kobayashi et al., 2014; Lawson and Weinstein, 2002; Lewis et al., 2004; Mahalwar et al., 2014; Moro et al., 2012; North et al., 2007). For simplicity in the text, these lines are referred to with short forms listed in square brackets: *Tg(kdrl:Cherry-CAAX) [kdrl:mCherry], Tg(fli1a:eGFP) [fli1a:eGFP], Tg(UAS:CA-β-catenin)^sd47Tg^ [UAS:CA-β-catenin], Tg(gata2b:KalTA4)[HSC:Gal4]* and *(UAS:Lifeact:eGFP) [UAS:eGFP], Tg(UAS:Cre) [UAS:Cre], Tg(bactin2:loxP-BFP-loxP-DsRed) [loxP BFP loxP dsRed]*.

MO for *fzd9b* was targeted to block the ATG start codon with sequence 5’-AGGTGAGCTTCCCATTCTGGATTTT-3’ from GeneTools. 1-cell stage zygotes were injected with 2ng of *fzd9b* MO and disruption of protein expression was confirmed with a fluorescently tagged mRNA. Suboptimal MO dosage was 0.5ng. The *wnt9a* MO has been previously described (Grainger et al., 2016b) and was used at 0.1ng. The *egfra* MO has been previously described (Zhao and Lin, 2013) and was used at 2.5ng or 1ng. Rescue experiments were performed using 20 pg *fzd* chimeric mRNA synthesized using the SP6 mMessage machine kit (Ambion) according to the manufacturer’s recommendations.

CRISPR/Cas9 was used to generate germline mutants for *fzd9b*; sgRNAs were chosen according to their ability to cleave DNA *in vitro* as previously described (Grainger et al., 2016a). Mutation of the *fzd9b* locus at the N-terminal was achieved 100ng of *cas9* mRNA (Trilink) and 100ng of sgRNA (GGCTCTTATGACCTGGAGAG) and generated mutants with either a 2bp insertion (*fzd9b^2bp^*) ^fb203^, or a 7bp deletion (*fzd9b^7bp^*)^fb204^. Similarly, mutation at the C-terminal tail was achieved by injecting the sgRNA (GGACTCTTCAGTGCCCACAG). For simplicity, in the text, these are referred to as *fzd9b^-/-^* and *fzd9b^ΔCTT/ΔCTT^*, respectively. Mutations were confirmed by sequencing individuals.

*Tg(fzd9b:Gal4)* founders were established by injecting 25pg of a Tol2 kit (Kwan et al., 2007) generated plasmid with 100pg of transposase mRNA at the 1-cell stage. The transgenic plasmid encoded a 4.3 kb *fzd9b* promoter region amplified using the primers: 5’ CTCCCATGAGGCAGAACGTGTGT 3’ and 5’ AGTCCGCGAGCAGCTTGTCTGTT 3’; this was cloned into a p5E MCS Tol2 entry vector using XhoI and SacII restriction sites and then combined with a Gal4 middle entry and polyA 3 prime entry vector by Gateway Assembly to make a full transgene construct with *cmlc2:gfp* in the backbone. The resultant animals were crossed to *Tg(UAS:YFP)*, and expression compared to *in situ* hybridization for *fzd9b* to identify founders that recapitulated endogenous *fzd9b* expression.

Lineage tracing experiments were visualized on a Zeiss LSM 880 with Airyscan. Representative images were produced by combining 3-4 Z-slices per scan.

### Whole-mount *in situ* hybridization (WISH) and Fluorescent WISH (FISH)

RNA probe synthesis was carried out according to the manufacturer’s recommendations using the DIG-RNA labeling kit, or the fluorescein labeling kit for FISH (Roche). Probes for *fli1a, rag1, dll4, dlc, notch1b*, *kdrl, cdh17, cmyb* and *runx1* and WISH protocols have been previously described (Clements et al., 2011; Kobayashi et al., 2014; Rowlinson and Gering, 2010). The probe for *fzd9b* was generated from the full-length cDNA. FISH signal was developed as previously described (Brend and Holley, 2009).

### Fluorescence activated cell sorting and quantitative PCR

Zebrafish were dissociated using Liberase TM (Roche) and filtered through an 80um filter. Cells were sorted on a BD Influx cell sorter according to standard procedures. RNA and cDNA were synthesized by standard means and qPCR was performed using FastStart Universal SYBR Green Master Mix (Roche) according to the manufacturer’s recommendations and analyzed using the 2^-∆∆Ct^ method (Schefe et al., 2006). Sequences of primers are shown in the table below.

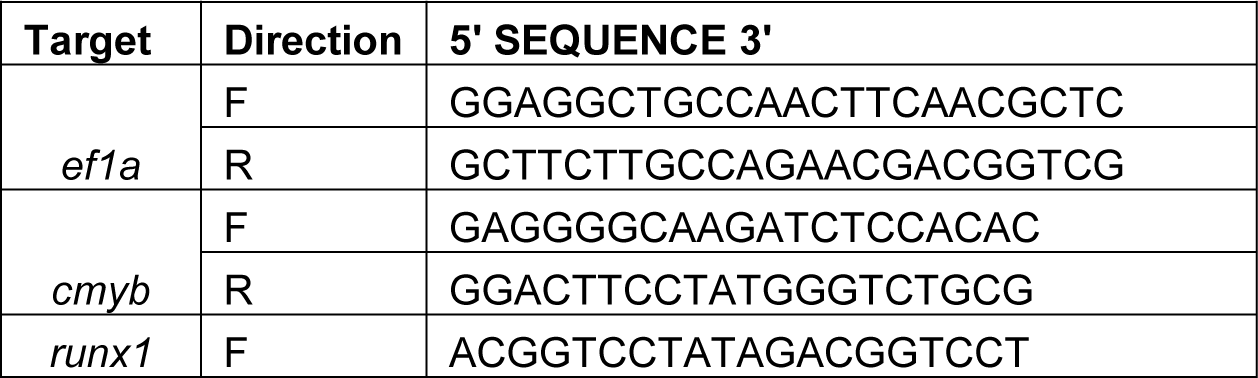
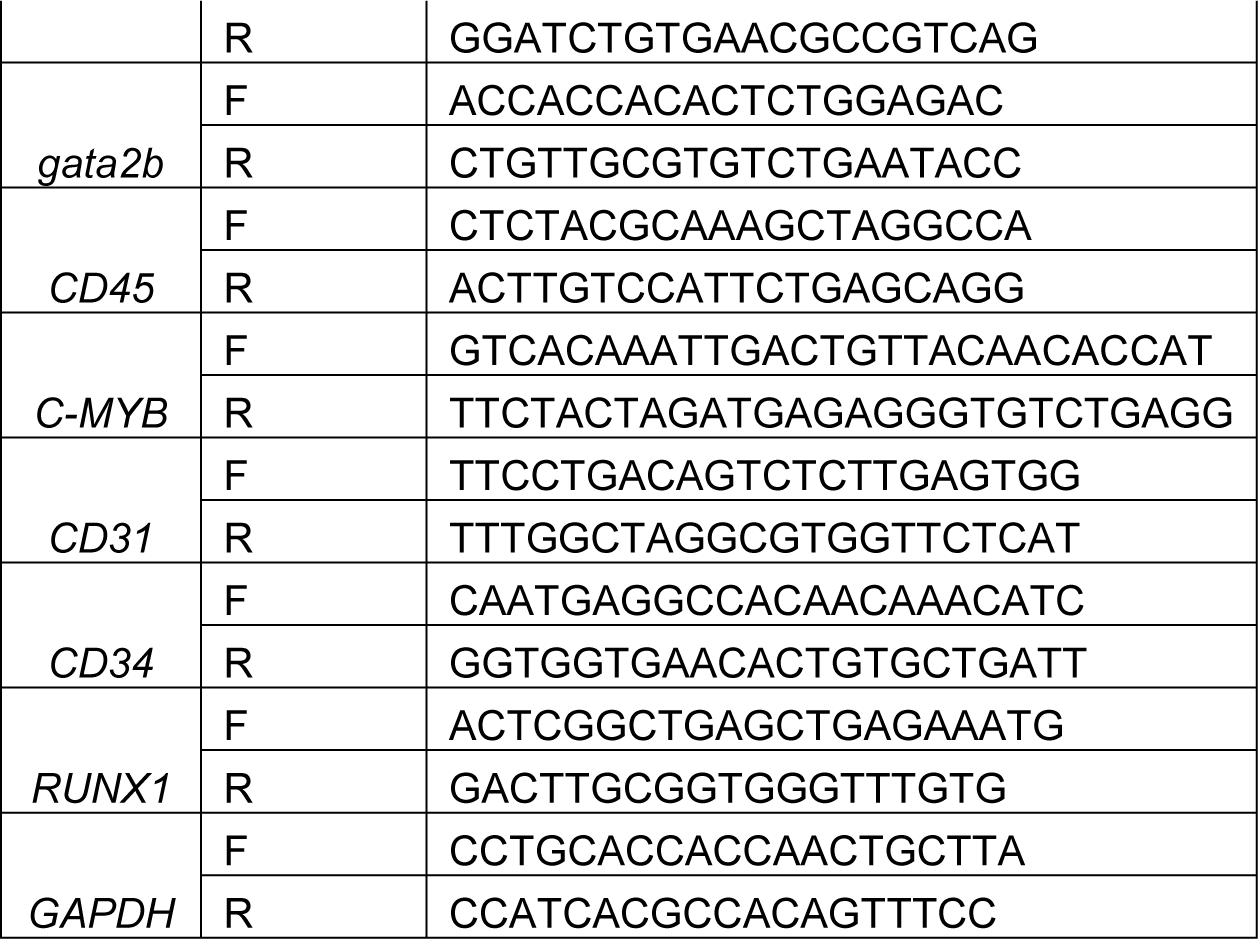

### Quantifying HSPCs

HSPCs were quantified by counting the number of *kdrl:mCherry; gata2b:GFP* double positive cells in floor of the DA in the region above the yolk extension in a 625μm confocal Z-stack encompassing the entire mediolateral segment of the aorta. The number of HSPCs per mm was calculated from this data. Confocal images were generated by stacking 1-4 individual Z-slices. When quantifying WISH data, the number of *cmyb*+ or *runx1+* cells were counted above the yolk extension.

### Human embryonic stem cell culture and HSPC differentiation

All experiments described in this study were approved by a research oversight committee (IRB/ESCRO Protocol #100210, PI Willert). Human embryonic stem (hES) cell H9 (WA09, NIH Registration number 062) cells were obtained from WiCell. Cells were maintained in Essential 8 (E8) media (Chen et al., 2011), with minor modifications, as previously described (Huggins et al., 2017). Plasmids encoding pools of shRNAs for FZD9 and WNT9A were obtains from ABM. The cell lines harboring shRNAs for control, WNT9A and FZD9 were generated by lentiviral transduction, as previously described (Moya et al., 2014). Virally-infected cells were selected with puromycin (4 ug/mL), and differentiated to HSPCs as previously described (Ng et al., 2008; Ng et al., 2005). Cells were dissociated and HSPCs were quantified by flow cytometry as previously described (Richter et al., 2018).

### APEX2-mediated proximity labeling

Chinese hamster ovary (CHO) cells were stably integrated with a CMV:Wnt9a construct. Conditioned medium from CHO cells with or without this construct was collected for two weeks, pooled together, filtered through a 0.22μm filter and tested for Wnt9a activity by luciferase STF assay with Fzd9b. HEK293T cells were stably integrated with a CMV:Fzd9b-5GS-APEX2 construct. APEX2-mediated proximity labeling was carried out essentially as described (Hung et al., 2016; Lam et al., 2015). Briefly, confluent 150mm plates of CMV:Fzd9b-5GS-APEX2 cells were treated with biotin-phenol for a total of 30 minutes each, ending at the time of hydrogen peroxide treatment. Cells were treated with Wnt9a conditioned medium for 1 or 5 minutes, or with WT CHO conditioned medium for 5 minutes. Cells were treated with hydrogen peroxide for 1 minute, quenched, lysed, and biotinylated proteins enriched by streptavidin pulldown, as previously described (Hung et al., 2016).

### Protein digestion

To denature the eluted proteins, an equal volume of 8 M Urea in 50 mM HEPES, pH 8.5 was added to each sample. Protein disulfide bonds were reduced by Dithiothreitol (DTT, Sigma) and alkylated with Iodoacetamide (Sigma) as previously described (Haas, 2006). Proteins were precipitated using trichloroacetic acid and resuspended in 300 μl of buffer (1 M urea (Fisher), 50 mM HEPES, pH 8.5) for proteolytic digestion.

Proteins were serially digested with 30 μg of LysC overnight at room temperature, then with 3 μg of trypsin for 6 hours at 37 °C (Lapek et al., 2017), quenched by the addition of trifluoracetic acid (TFA, Pierce), and peptides were desalted with C18-StageTips extraction columns (Waters) as previously described (Lapek et al., 2017). Peptides were dried in a speed vac, then re-suspended in 50% Acetonitrile/5% formic acid and quantified by the Pierce Quantitative Colorimetric Peptide Assay (Thermo); an equal amount of each sample to be run on a pooled bridge channel (Lapek et al., 2018). Aliquots were dried under vacuum and stored at −80°C until they were labeled with Tandem Mass Tag (TMT) reagents.

### TMT labeling

Peptides were labeled with 10-plex TMT reagents (Thermo) (McAlister et al., 2012; Thompson et al., 2003) as previously described (Lapek et al., 2017). Briefly, TMT reagents were reconstituted in dry acetonitrile (Sigma) at 20 μg/mL. Dried peptides were re-suspended in 30% dry acetonitrile in 200 mM HEPES, pH 8.5, and 8 μL of the appropriate TMT reagent was added to the peptides. Reagent 126 (Thermo) was used to bridge between mass spectrometry runs. Remaining reagents were used to label samples in a random order. Labeling was carried out for 1 hour at room temperature and was quenched by adding 9 μL of 5% hydroxylamine (Sigma) which was allowed to react for 15 mins at room temperature. Labeled samples were acidified by adding 50 μL of 1%TFA, pooled into appropriate 10-plex TMT samples and desalted with C18 Sep-Paks.

### LC-MS2/MS3 Analysis

All LC-MS2/MS3 experiments were performed on an Orbitrap Fusion mass spectrometer (Thermo) with an in-line Easy-nLC 1000 (Thermo). Home-pulled, home-packed columns (100 mm ID x 30 cm, 360 mm OD) were used for analysis. Analytical columns were triple-packed with 5 μm C4 resin, 3 μm C18 resin, and 1.8 μm C18 resin (Sepax) to lengths of 0.5 cm, 0.5 cm, and 30 cm, respectively. Peptides were eluted with a linear gradient from 11 to 30% acetonitrile in 0.125% formic acid over 165 min at a flow rate of 300 nL/minute and heating the column to 60°C. Nano-electrospray ionization was performed by applying 2000 V through a stainless-steel T-junction at the inlet of the microcapillary column.

The mass spectrometer was operated in a data-dependent mode, with a survey scan performed over a mass to charge (m/z) range of 500–1200 at a resolution of 1.2 x 10^5^ in the Orbitrap. The target automatic gain control (AGC) was set to 2 x 10^5^ with a maximum inject time of 100 ms and an s-lens RF of 60. Top Speed mode was used to select the most abundant ions for tandem MS analysis. All data collected was centroided.

Ions above an intensity threshold of 5 x 10^5^ were isolated in the quadrupole and fragmented using collision-induced dissociation (normalized energy: 30%) for MS2 analysis. MS2 fragments were detected in the ion trap using the rapid scan rate setting with an AGC of 1 x 10^4^ and a maximum injection time of 35 ms.

For MS3 analysis, synchronous precursor selection was used to maximize quantitation sensitivity of the TMT reporter ions (McAlister et al., 2014). Up to 10 MS2 ions were simultaneously isolated and fragmented with high energy collision induced dissociation (normalized energy: 50%). MS3 fragment ions were analyzed in the Oribtrap with a resolution of 6 x 10^4^. The AGC was set to 5 x 10^4^ using a maximum injection time of 150 ms. MS2 ions 40 m/z below and 15 m/z above the MS1 precursor ion were excluded from MS3 selection.

### Data Processing

Raw spectral data were processed using Proteome Discoverer 2.1.0.81 (Thermo). MS2 spectra were identified using the Sequest HT node (Eng et al., 1994), searching against the Human Uniprot database (downloaded: 5/11/2017) with the zebrafish Fzd9b sequence appended. False discovery rate (FDR) estimation was performed using a reverse decoy database (Elias and Gygi, 2007; Elias et al., 2005; Peng et al., 2003). Search parameters were as follows. Mass tolerances were set to 50 ppm and 0.6 Da for MS1 and MS2 scans, respectively. Full trypsin digestion was specified with a maximum of two missed cleavages per peptide. Modifications included static 10-plex TMT tags on peptide n-termini and lysine, static carbamidomethylation of cysteine and variable oxidation of methionine. Data were filtered to a 1% FDR at both the peptide and protein level.

The intensities of TMT reporter ions were extracted from the MS3 scans for quantitative analysis. Prior to quantitation, spectra were filtered to have an average signal to noise of 10 across all labels and an isolation interference less than 25%. Data were normalized in a two-step process as previously described (Lapek et al., 2018), by normalizing each protein the pooled bridge channel value and then normalizing to the median of each reporter ion channel and the entire dataset. The dataset has been uploaded to ProteomeXchange (PXD010649) through MassIVE (MSV000082677).

### Statistical Analysis

For APEX results, two-tailed student’s t-tests were used to determine significantly enriched proteins at each time point. If the variances between samples were determined to be unequal by an F-test, Welch’s correction was used. Significantly changing proteins were prioritized using pi score (Xiao et al., 2014), a metric that takes both p-value and fold-change into account. Gene ontology of the significant proteins was performed using the database for annotation, visualization and integrated discovery (DAVID) server (Huang da et al., 2009; Huang et al., 2007). For STF assays and qPCR or cell counting comparing more than two populations, one-way ANOVA, followed by Bonferroni post-test analysis were conducted. For qPCR or cell counting comparing only two populations, two-tailed student’s t-tests were used.

### Plasmids

Expression constructs were generated by standard means using PCR from cDNA libraries generated from zebrafish larvae at 24 hpf, or from hES cells; these were cloned into pCS2+, downstream of a CMV promoter, and upstream of *IRES:mKate2*. Addgene provided expression vectors for *Cas9* (47929), guide RNAs (46759) and zebrafish *ctnnb1* (17199).

## Author Contributions

SG conceived, designed and conducted experiments and analysis, and wrote the manuscript. NN and JR designed and conducted experiments and analysis, JS, BL, RB, CHO and JH conducted experiments and analysis, JMW, MCT and DG performed mass spectrometry and analysis, CK and ID provided zebrafish lines, DT and KW supervised experiments and edited the manuscript.

## Acknowledgements

C. Fine and J. Olvera for FACS assistance; C. Pouget and P. Sahai-Hernandez for technical assistance and critical reading of the manuscript; R. Espin-Palazon, J. Nussbacher, I.J. Huggins and N. Grimsey for critical reading of the manuscript; I.J. Huggins for antibodies; N.Gohad for microscopy assistance. SG was supported by awards from the American Heart Association (14POST18340021), the Leukemia and Lymphoma Society (5431-15) and the NIH/NHLBI (K99HL133458). Research reported in this publication was supported by the National Heart, Lung, And Blood Institute of the National Institutes of Health under Award Number K99HL133458. The content is solely the responsibility of the authors and does not necessarily represent the official views of the National Institutes of Health. JR was supported by was supported in part by the UCSD Interdisciplinary Stem Cell Training Program (CIRM TG2-01154) and by the American Heart Association Predoctoral Fellowship 16PRE27340012. This work was supported in part by funding to KW from the UCSD Stem Cell Program and was made possible in part by the CIRM Major Facilities grant (FA1-00607) to the Sanford Consortium for Regenerative Medicine. DT was supported by Scholar Award (1657-13) from The Leukemia and Lymphoma Society and CIRM (RB4-06158). DT and KW were supported in part by NIH/NHLBI under the Grant R01HL135205. JMW was supported by Graduate Training in Cellular and Molecular Pharmacology Training Grant NIH T32 GM007752. Harvard Stem Cell Institute and NIDDK grant UH2/3DK107372 to I.A.D.

